# Synaptic weights that correlate with presynaptic selectivity increase decoding performance

**DOI:** 10.1101/2022.11.28.518203

**Authors:** Júlia V. Gallinaro, Benjamin Scholl, Claudia Clopath

**Affiliations:** Bioengineering Department, Imperial College London, London, UK; Department of Neuroscience, Perelman School of Medicine, University of Pennsylvania, Philadephia, PA, USA

## Abstract

The activity of neurons in the visual cortex is often characterized by tuning curves, which are thought to be shaped by Hebbian plasticity during development and sensory experience. This leads to the prediction that neural circuits should be organized such that neurons with similar functional preference are connected with stronger weights. In support of this idea, previous experimental and theoretical work have provided evidence for a model of the visual cortex characterized by such functional subnetworks. A recent experimental study, however, have found that the postsynaptic preferred stimulus was defined by the total number of spines activated by a given stimulus and independent of their individual strength. While this result might seem to contradict previous literature, there are many factors that define how a given synaptic input influences postsynaptic selectivity. Here, we designed a computational model in which postsynaptic functional preference is defined by the number of inputs activated by a given stimulus. Using a plasticity rule where synaptic weights tend to correlate with presynaptic selectivity, and is independent of functional-similarity between pre- and postsynaptic activity, we find that this model can be used to decode presented stimuli in a manner that is comparable to maximum likelihood inference.

## Introduction

Neurons in the visual cortex are selectively driven by specific features of sensory stimuli. This selective neural activity is proposed to be shaped by Hebbian plasticity (Hebb 1949) during developmental stages and sensory experience. Hebbian plasticity can be described as the strengthening of synaptic weights in a manner dependent on the correlation of the pre- and postsynaptic neurons (Gerstner and Werner M. 2002; Gerstner and Kistler 2002). Thus, in the development of visual cortical circuits, Hebbian plasticity is thought to lead to a functional distribution in synaptic weights: weights are larger between pre- and postsynaptic neurons with similar functional preferences (Clopath et al. 2010) and only a few synaptic inputs are needed to define postsynaptic sensory feature selectivity (Goetz et al. 2021). In support of this framework, excitatory pyramidal neurons in layers 2/3 of mice primary visual cortex are shown to form functional subnetworks; the synaptic weights between neurons and the probability of connectivity between neurons reflect the similarity in tuning to visual features, such as for example orientation preference (Harris and Mrsic-Flogel 2013; Ko et al. 2011; Ko et al. 2013; Cossell et al. 2015; Lee et al. 2016).

However, a recent study (Scholl et al. 2021) has shown different results in the visual cortex of ferrets. Instead of a functionally-defined weight distribution, Scholl et al. 2021 found that preference for orientation stimuli is derived from the total number of excitatory synaptic inputs activated by a given stimulus. That is, strong and weak synapses were recruited for all visual stimuli presented. While this result appears to contradict the previous literature, there are many factors which impact how a given synaptic input might influence activity at the somatic output and postsynaptic selectivity. Some of these factors include synapse weight and number, reliability (Branco et al. 2008), location within the dendritic tree (Stuart and Spruston 1998), co-activity with their neighbors (Scholl et al. 2017), and presence of dendritic inhibition (Gidon and Segev 2012). In fact, given that pairs of cortical neurons tend to be connected through multiple synaptic contacts (Feldmeyer et al. 1999; Feldmeyer et al. 2002; Feldmeyer et al. 2006; Markram et al. 1997), an overall stronger weight between pairs of pre- and postsynaptic with correlated activity could be achieved through many contacts with a mixture of weights. In this case, weights might encode other features of presynaptic activity. For example, Scholl et al. 2021 also showed that anatomical correlates of synaptic strength correlate with spine selectivity, i.e. the sharpness of the tuning curve or how selectively a spine responds to specific orientations.

Here, we explore how synaptic weights based on presynaptic selectivity, rather than functional-similarity between presynaptic and postsynaptic activity, might be achieved by cortical neurons and what impacts this distribution has on neuron decoding. We model a feedforward circuit with a plasticity rule leading to synaptic weights that are correlated with presynaptic selectivity (termed ‘variance rule’), independent of the difference between pre- and postsynaptic preferred orientation (ΔPO). We show that this plasticity rule can lead to postsynaptic neurons selectively tuned to the specific stimuli. Furthermore, we show that a decoder which uses the weights emerging from the plasticity rule performs comparatively to a decoder derived based on maximum likelihood inference. Overall, our results suggest a decoding model where the somatic preference is defined mostly by the number of activated spines given a certain stimulus, while the strength of individual spines reflects input selectivity.

## Results

Here we study the functional implications of a plasticity rule based on presynaptic variance, rather than covariance between presynaptic and postsynaptic activity (i.e. classic Hebbian), in a feedforward circuit. Our circuit is composed of one postsynaptic neuron, modeled as a point neuron (see Methods for details), which receives input from *N* presynaptic neurons through plastic weights *w_i_*, and from an untuned inhibitory source (Figure 1A). Presynaptic neurons are orientation selective and their activity is modulated by tuning curves (Figure 1B). In order to model a diverse population (i.e. differences in orientation preference and selectivity), the tuning curve of each presynaptic neuron *i* has its own preferred orientation (PO, 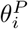) and width (*κ_i_*) (Figure 1B).

**Figure 1:**
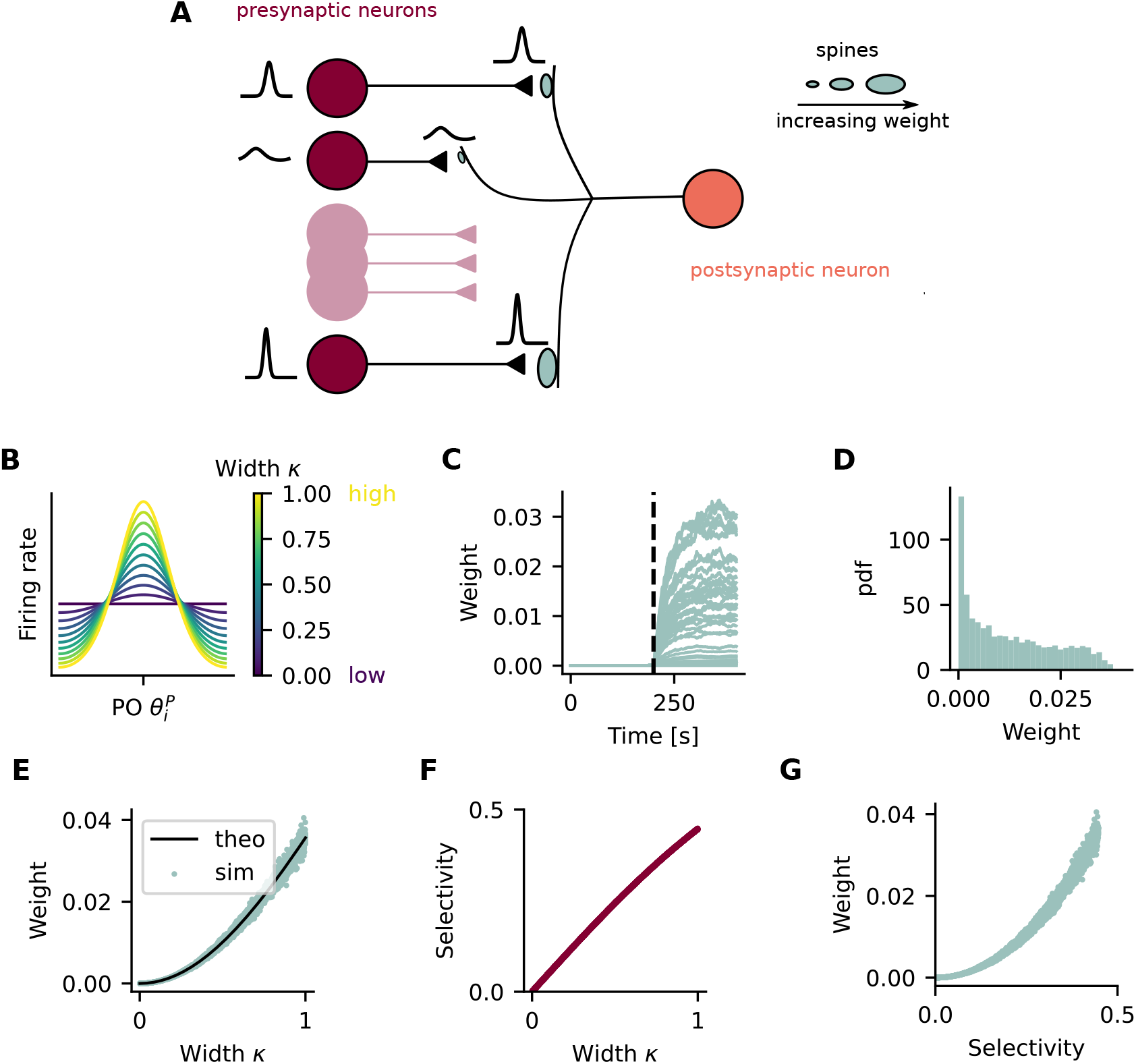
A plasticity rule based on presynaptic variance results in weights that correlate with presynaptic selectivity. (A) A postsynaptic point neuron receives input from 50 presynaptic neurons. Synaptic weights, represented by the spine sizes, are plastic according to a rule based on presynaptic variance. (B) The activity of presynaptic neurons is moduled by von Mises tuning curves. The tuning of the presynaptic neuron *i* is defined by its PO 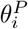 and its selectivity, given by the width parameter *κ_i_*. (C) Synaptic weights as a function of time. Each line shows the synaptic weight of a single presynaptic neuron. The vertical dotted line indicates the moment when the stimulation protocol starts. (D) Distribution of synaptic weights at the end of the stimulation protocol. (E) Synaptic weights as a function of the width parameter *κ*. Green dots show data obtained from simulation and black line from theory. (F) Selectivity of presynaptic neurons as a function of their width parameter *κ*. (G) Synaptic weight at the end of simulation as a function of presynaptic selectivity. (D-G) Shown are data pooled from 100 independent simulation runs.

### A plasticity rule based on presynaptic variance results in weights that correlate with presynaptic selectivity

In order to build a model where synaptic weights correlate with presynaptic selectivity, we first study how synaptic plasticity might lead to this correlation. Based on previous studies showing that some forms of LTP can be induced without the need of postsynaptic depolarization (Urban and Barrionuevo 1996; Ito and Sugiyama 1991; Katsuki et al. 1991; Zalutsky and Nicoll 1990), we propose a plasticity rule in which potentiation is based on presynaptic activity only. More specifically, a rule in which potentiation is based on presynaptic variance (variance rule). In this way, changes in synaptic weight, Δ*w*, can be summarized as

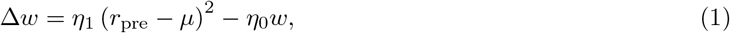

where *r_pre_* is the presynaptic activity, *μ* is a constant representing the mean presynaptic activity, *η*_1_ is the learning rate and *η*_0_ is the weight decay rate. We then stimulate the presynaptic neurons by showing a stimulus of orientation *θ*, which changes every 200 ms, and observe the evolution of the synaptic weights (Figure 1C).

Once converged, the synaptic weights have an asymmetric distribution (Figure 1D), and are essentially a function of the width of the presynaptic tuning curve (Figure 1E, see also). Since selectivity of a presynaptic neuron is given by the width of its tuning curve *κ* (Figure 1F), the converged weights are also correlated with presynaptic selectivity (Figure 1G). Thus, we show that a feedforward model with a plasticity rule based on the variance of presynaptic activity, leads to synaptic weights that are correlated with presynaptic selectivity.

### Postsynaptic neurons are orientation selective and synaptic weights are uncorrelated with the difference in preferred orientation between pre- and postsynaptic neuron (ΔPO)

In the previous simulations, we find that, similar to the presynaptic neurons, the postsynaptic neuron is also orientation tuned (Figure 2B). Extracting the relevant parameters from the postsynaptic tuning curve across multiple simulation runs shows that individual postsynaptic neurons respond preferentially to orientations spread across the full range of stimuli 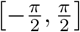 and with selectivity values that cover almost the entire range [0, 1] (Figure 2C). Note that the value of selectivity depends on the mean activity level (Merkt et al. 2019), which in our simulations is strongly influenced by the amount of inhibition received by the postsynaptic neuron (Supplementary Figure 1).

**Figure 2:**
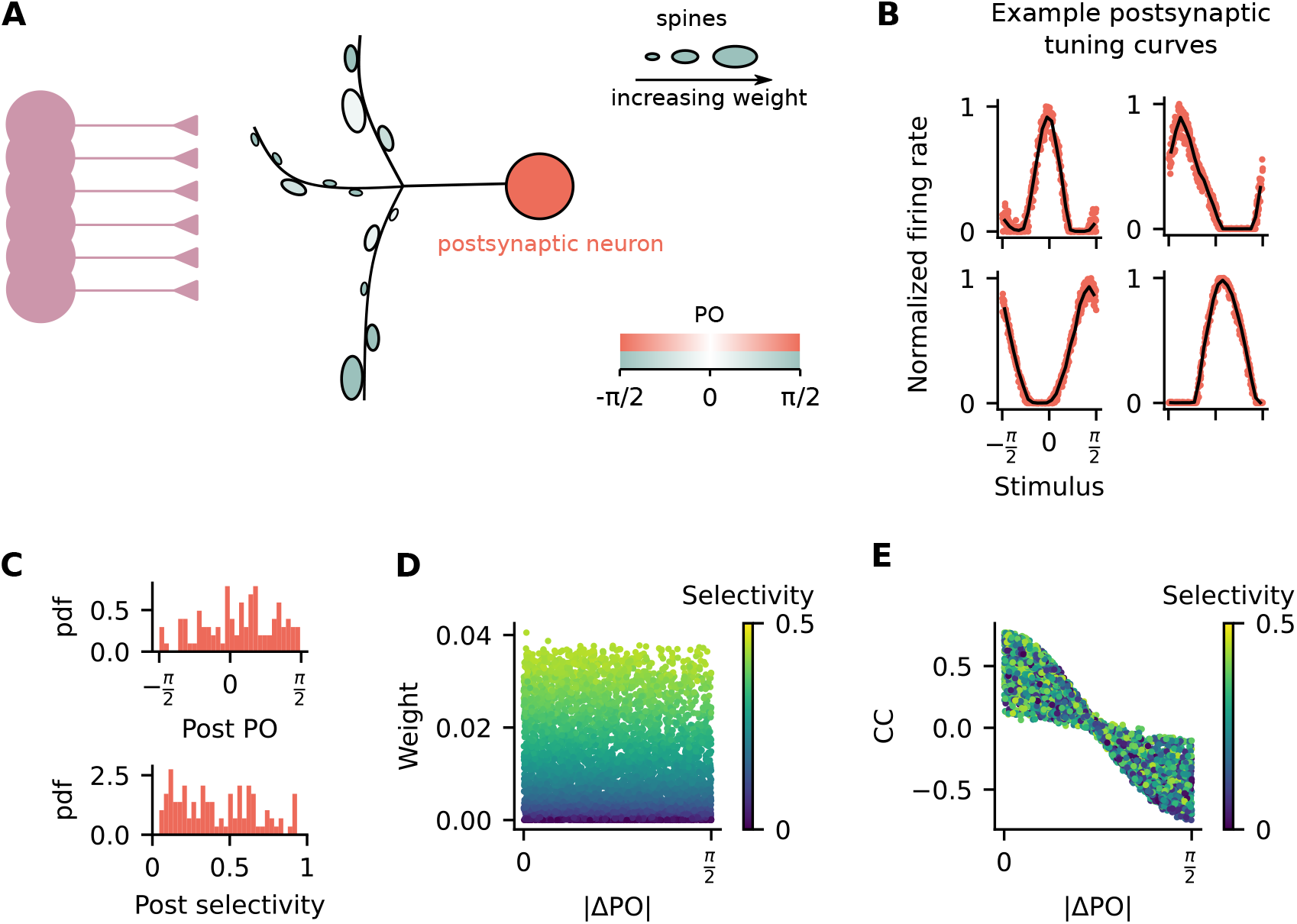
Postsynaptic neurons are orientation selective and synaptic weights are uncorrelated with the difference in preferred orientation between pre- and postsynaptic neuron (ΔPO). (A) A postsynaptic neuron receives input from 50 presynaptic neurons. Spine sizes represent synaptic weights and spine colors represent PO of the presynaptic neurons. The postsynaptic neuron is orientation selective and its PO is represented by its color. (B) Examples of tuning curves for the postsynaptic neuron from 4 independent simulation runs. Firing rates are normalized by the maximum rate of each individual tuning curve. (C) Statistics of postsynaptic selectivity. Data pooled from 100 independent simulation runs. *Top*: Probability density function (pdf) of the postsynaptic PO. *Bottom*: Probability density function (pdf) of the postsynaptic selectivity. (D) Synaptic weights as a function of ΔPO. (E) Pearson’s correlation coefficient between pre- and postsynaptic neurons as a function of ΔPO. (D-E) Colors indicate presynaptic selectivity.

Synaptic plasticity rules which are based on the covariance between pre- and postsynaptic activity generate circuits in which the strength of synaptic weights are anti-correlated with ΔPO (Supplementary Figure 2). That is, if the pre- and postsynaptic neurons have similar tuning, they typically are connected with a strong weight. But this is not necessarily the case for a rule based on the variance of presynaptic activity only. Therefore, we next test whether the synaptic weights are anti-correlated with ΔPO. Similar to experimental data (Scholl et al. 2021), we find that the synaptic weights are not correlated with ΔPO and depend mostly on presynaptic selectivity (Figure 2D). Even though there is no clear relationship between the synaptic weights and ΔPO, the activity of the postsynaptic neuron is still more correlated with the activity of presynaptic neurons that have similar POs (Figure 2E).

### Postsynaptic neurons inherit preferred orientation from the number of presynaptic inputs with similar preference

If it is not true that synaptic weights are stronger when pre- and postsynaptic neurons have similar POs, what is then defining the postsynaptic orientation preference? While some previous work have proposed that somatic functional preference is derived from the functional preference of a few stronger presynaptic inputs (Cossell et al. 2015; Lee et al. 2016), others have proposed that it is actually defined by the number of active spines with a given PO (Scholl et al. 2021). Since in our model the synaptic weights are not correlated with ΔPO (Figure 2D), we expect that somatic preference will be defined by the number of presynaptic inputs with similar preferences.

In order to test this, we wanted to know whether all orientations within the interval 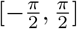 were equally represented within presyanptic populations or if there was any bias towards the PO of the postsynaptic neuron. Even though the PO of the presynaptic inputs were independently drawn from a uniform distribution (Figure 3A), we find that there are indeed slightly more presynaptic inputs which are co-tuned than inputs which are not co-tuned with the postsynaptic neuron (Figure 3B). To further explore this result, we calculate the total input current to the postsynaptic neuron when different orientations are being shown to the presynaptic population (Figure 3C). We then split the total input current into two values: the total number of active inputs and the mean synaptic weight of those active inputs. Inputs are considered to be active when their firing rate is above a certain threshold during the presentation of a stimulus (see Methods for details). We find that, similar to the total input current (Figure 3C), the number of active inputs is also modulated by the shown stimulus such that a larger number is active when the PO is shown (Figure 3D).

**Figure 3:**
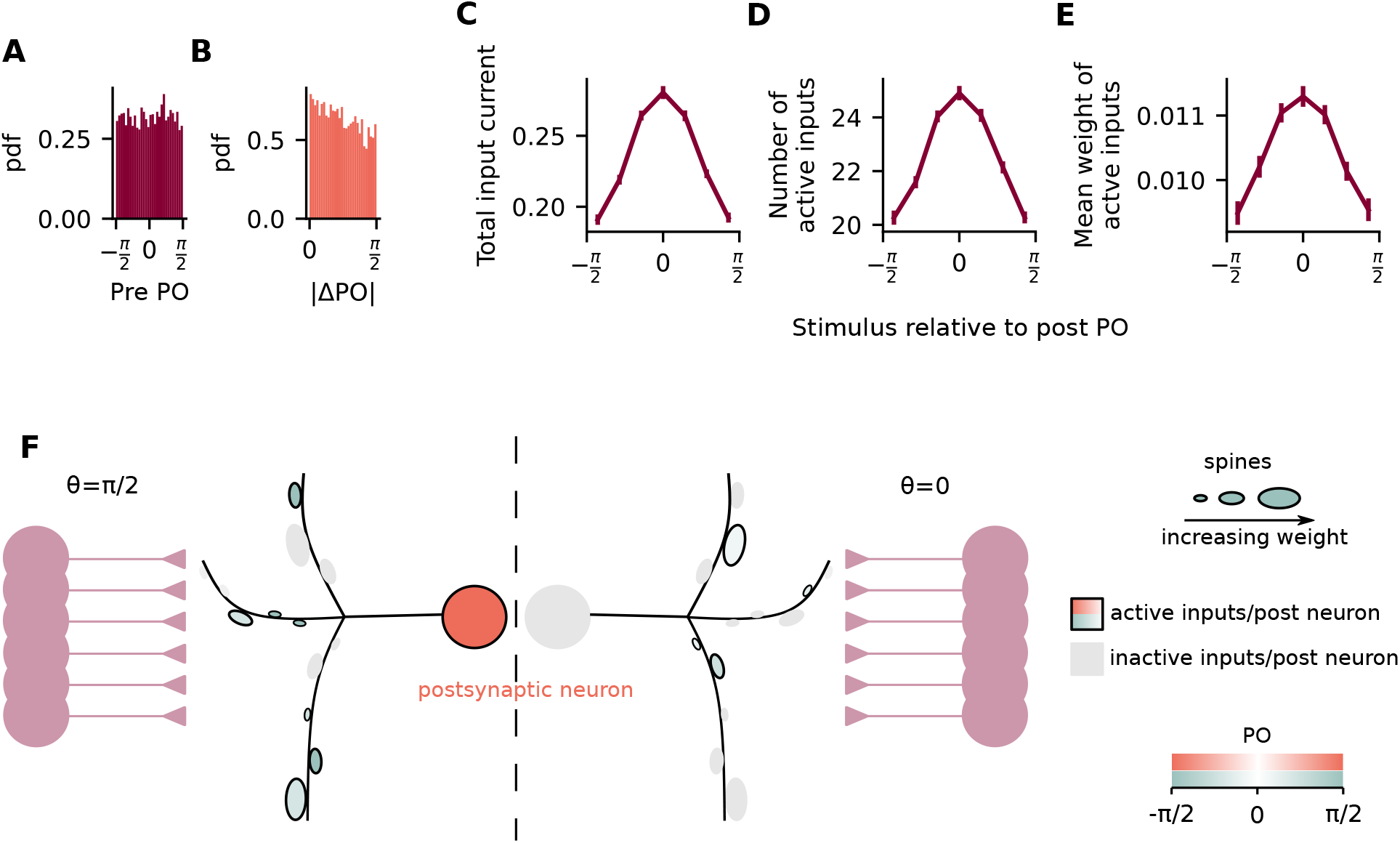
Postsynaptic neurons inherit preferred orientation from the number of presynaptic inputs with similar preference. (A) Histogram of the PO of presynaptic neurons. (B) Histogram of the difference in PO between pre- and postsynaptic neurons. (C) Total input current to postsynaptic neuron when different orientations are being shown. (D) Number of active inputs when different orientations are being shown. (E) Mean weight of active inputs when different orientations are being shown. (F) A postsynaptic neuron receives inputs from 50 presynaptic neurons. When an orientation is shown to the presynaptic neurons, only a fraction of them have their activity above a certain threshold (active inputs, represented as colored spines), while the rest have their activity below the threshold (inactive inputs, represented as grey spines). When the PO of the postsynaptic neuron is shown (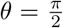, *left*), there are on average more active inputs than when the orthogonal orientation is shown (*θ* = 0, *right*).

Unexpectedly, however, we find that the mean weight of the active inputs is also modulated by the difference between the shown stimulus and the postsynaptic preferred stimulus (Figure 3E). This is different to what was observed in experiments (Scholl et al. 2021) and counter-intuitive given that the plasticity rule used here is based on presynaptic activity only. However, even though correlated activity between pre- and postsynaptic neurons does not lead directly to stronger weights, the postsynaptic PO is given by a sum of the presynaptic POs weighted by their selectivity and by their input weights (Supplementary Figure 3). Therefore, the postsynaptic PO will be biased by presynaptic neurons with stronger weights, which are also the ones with higher selectivity (Supplementary Figure 3).

### A decoder based on the variance rule performs comparatively to a decoder based on maximum likelihood

Hebbian plasticity has been previously shown to enhance orientation selectivity (Sadeh et al. 2015), in accordance with experimental studies showing an increase in orientation selectivity after visual experience (Ko et al. 2013; Hoy and Niell 2015). Since Hebbian plasticity shapes circuits in which synaptic weights are anti-correlated with ΔPO (Supplementary Figure 2), the question emerges of whether there is any computational advantage of synaptic weights that correlate with presynaptic selectivity instead. Therefore we next study a feedforward circuit where the synaptic weights are correlated with presynaptic selectivity in the context of decoding stimuli.

We assume there is a population of input neurons with different tuning curves (Figure 4A). The same orientation *θ* is shown to all input neurons and the firing rate of presynaptic neuron i is given by the shown stimulus and their respective tuning curve, defined by their PO 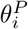 and width *κ_i_*. A decoder receives inputs from all neurons in this population and estimates the shown orientation based on their tuning curves and firing rates (Yates and Scholl 2022). We then derive a decoder, which is based on maximum likelihood inference (ML decoder, see Methods for details) (Abbott and Dayan 2005). Assuming the input tuning curves to be modeled as von Mises functions, the ML decoder estimates the shown orientation with (Figure 4B):

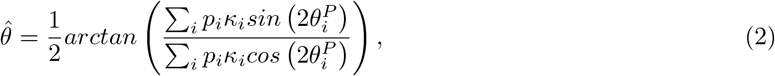

where *p_i_* is the firing rate of neuron *i* (*r_i_* with added Poisson noise, see Methods for details), *κ_i_* is a parameter of the von Mises tuning curve defining its width, 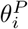 is the preferred stimulus of input neuron *i* and 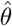 is the orientation estimated by the ML decoder.

**Figure 4:**
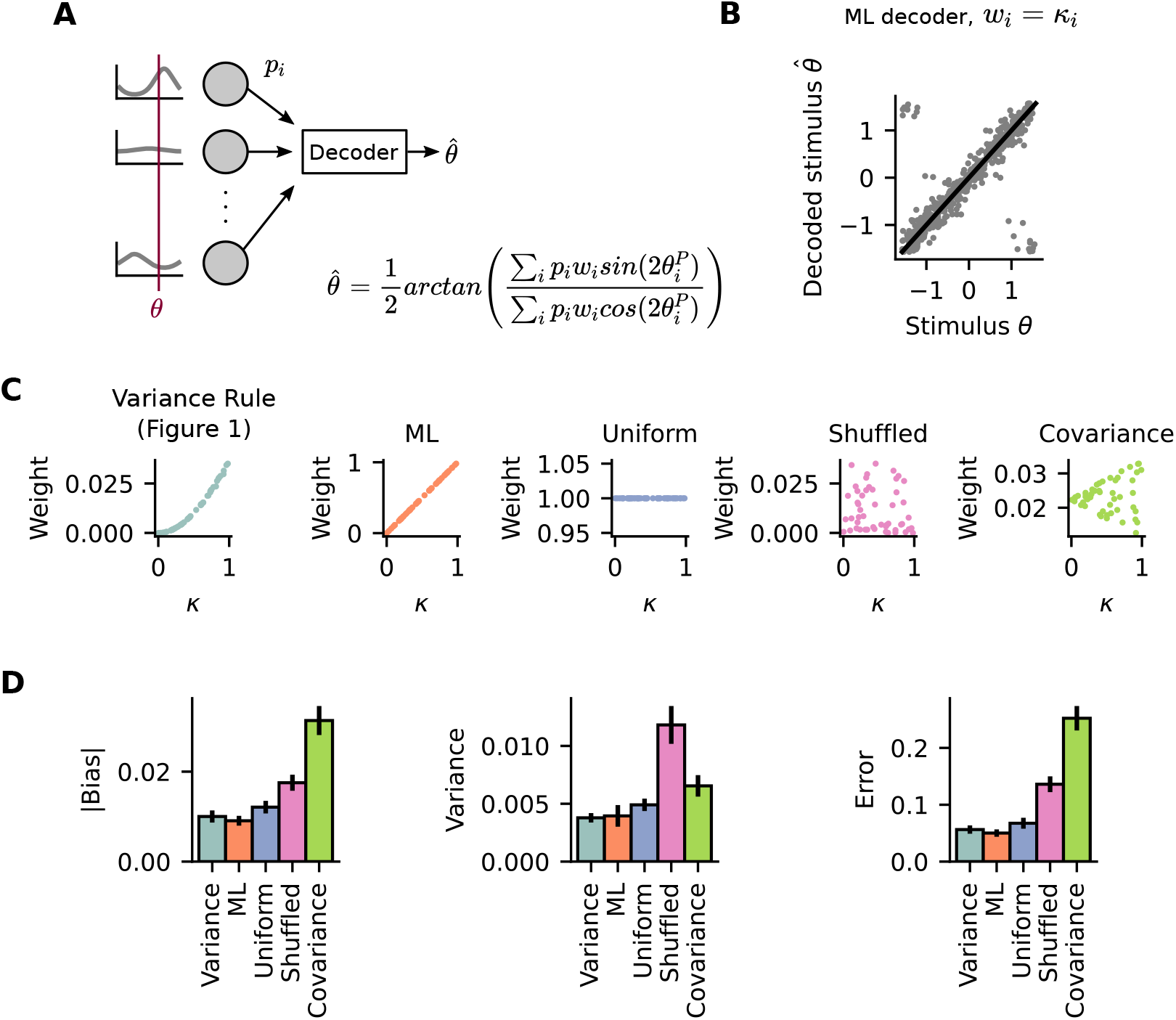
A decoder based on the variance rule performs comparatively to the ML decoder. (A) A decoder receives input from 50 input neurons and decodes the orientation that was shown to the whole population using the presented formula. (B) Decoded orientation 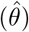 plotted against the actual orientation shown to the input population (*θ*) for a decoder based on maximum likelihood inference (ML decoder, *w_i_* = *κ_i_*). (C) Weight as a function of the presynaptic width parameter *κ* for five different decoders. (D) Performance of 5 decoders. *left*: Absolute value of bias, *middle*: variance and *right*: error. Bars show mean and standard error of the mean across 100 independent simulation runs.

Similar to Bayesian cue integration, in which inputs are integrated by being multiplied by their uncertainty (Knill and Pouget 2004; Ma et al. 2006; Echeveste and Lengyel 2018), this decoder is such that the firing rate *p_i_* of input neuron *i* is being multiplied by the width of its tuning curve *κ_i_*, which defines its selectivity (Equation 2). This multiplication *p_i_κ_i_* could be easily implemented by synaptic weights which are correlated with presynaptic selectivity (Figure 1). Therefore, we next compare the performance of the ML decoder with that of a decoder that uses the weights obtained from the variance rule (Figure 1). In order to do this, we assume that the decoder multiplies the firing rate of each presynaptic neuron *i* by a weight *w_i_*:

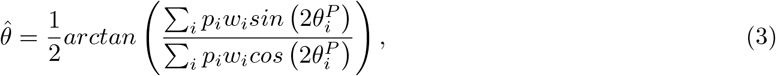

where *w_i_* are the weights used by the decoder and *w_i_* = *κ_i_* in the case of the ML decoder.

We then compare the performance of the decoder that uses the weights obtained from the variance rule (Variance Rule) to other decoders (Figure 4C), namely: (i) to a decoder where *w_i_* = *κ_i_* (ML); (ii) to a decoder that uses the same weight for all input neurons *w_i_* = 1 (Uniform); (iii) to a decoder that uses the same weights as the Variance Rule decoder but shuffled, and therefore uncorrelated to presynaptic selectivity (Shuffled); (iv) to a decoder that uses the weights obtained from a simulation using a covariance plasticity rule (Covariance, see 2). We find that the presynaptic variance decoder performs comparatively to the ML decoder and better than the others (Figure 4D).

In conclusion, we demonstrate how a plasticity rule based on presynaptic variance can lead to the formation of feedforward circuits where the postsynaptic neurons are selective for specific stimuli. Under this framework, we find that postsynaptic neurons can act as decoders whose performance rivals that of maximum likelihood inference. These results suggest a decoding model where individual inputs received by the post-synaptic neuron are weighted according to presynaptic selectivity rather than functional-similarity between pre- and postsynaptic activity.

## Discussion

In this work, we simulated a feedforward network in which weights were changing according to a plasticity rule based on the variance of presynaptic activity. The presynaptic population was composed of neurons with diverse PO and selectivity. We showed that, by stimulating the input population with different orientations, synaptic weights converged to values that were a function of presynaptic selectivity. We then showed that a decoder which used those weights to decode a stimulus had a performance comparable to a decoder based on maximum likelihood inference.

In the current paper, we focused on the computational implications of synaptic weights that correlate with presynaptic selectivity. We did not explore, however, the functional advantages of somatic preference being defined by the number of spines rather than by their strength (Scholl et al. 2021). This could be interesting to explore in view of recent experimental evidence showing that learning is associated with the formation of new spines (Hedrick et al. 2022; Fu et al. 2012; Hofer et al. 2009; Xu et al. 2009) and previous theoretical work suggesting that having multiple synaptic contacts between the same pair of neurons, instead of having a single strong one, could lead to more robust circuits with stable memories (Deger et al. 2018; M. Fauth et al. 2015; M. J. Fauth and Rossum 2019). A possible extension to our model, therefore, could be to include structural plasticity such that multiple contacts between pre- and postsynaptic neurons would be allowed to form.

Different structural plasticity rules have been proposed to describe activity dependent spine turnover (M. Fauth and Tetzlaff 2016). In a future study, using a structural plasticity rule forming stronger connectivity between neurons with similar orientation preference (Gallinaro and Rotter 2018), for example, could lead to a circuit where more synaptic contacts are formed between the same pair of pre- and postsynaptic neurons when their activity is correlated (Figure 2E). As a consequence, there could be an even stronger bias in the number of presynaptic inputs that are co-tuned with the soma (larger number of inputs with |ΔPO| = 0 in Figure 3B). Another possible consequence could be a stronger overall synaptic weight - i.e. synaptic weight of individual spines summed across all synaptic contacts - between pairs of co-tuned pre- and postsynaptic neurons. This could be interesting when comparing experimental data derived from measuring synaptic weights as anatomical features of spines versus measuring synaptic weights as amplitudes of postsynaptic potentials. On the other hand, using a structural plasticity rule in which weaker weights are more likely to be deleted than stronger ones (M. Fauth et al. 2015; Le Bé and Markram 2006) could lead to different functional circuits.

Previous work has shown that the diversity of synaptic weights found in the visual cortex of ferrets could be explained by a model where a decoder reads information from a population of diverse PO and selectivity (Yates and Scholl 2021). In a similar scenario, here we propose that the synaptic weights should reflect presynaptic selectivity. Such diversity in synaptic weights, therefore, would not be necessary if all presynaptic inputs had the same selectivity. Why then would there be a population of neurons with diverse selectivity in the visual cortex? One possibility is that decoding from a population of input neurons with a range of selectivity provides better discrimination capabilities for natural images (Goris et al. 2015). Another option, however, is that selectivity could somehow encode presynaptic uncertainty, but how exactly uncertainty is encoded in neural representation is still unclear (Koblinger et al. 2021).

In our model, if presynaptic selectivity would reflect input uncertainty, there would be the following consequences. Firstly, according to the rule presented in this paper, synaptic weights would fluctuate as the selectivity changes. Assuming that selectivity would reflect uncertainty, synaptic weight fluctuations would also reflect presynaptic uncertainty. This is similar to what has been recently proposed in a theoretical paper (Aitchison et al. 2021), and in accordance with experimental data showing that synaptic weights fluctuate over time (Hazan and Ziv 2020; Ziv and Brenner 2018). In contrast to our model, however, these experimental studies have shown that the weights can also fluctuate in an activity independent manner. Secondly, as the synaptic weights change according to presynaptic selectivity, or uncertainty, so would the postsynaptic PO. The postsynaptic PO would therefore change over time, reflecting the reliability of presynaptic inputs.

In conclusion, our results suggest a model of the visual cortex in which postsynaptic preferred orientation is defined by the number of spines with a given preferred orientation, while synaptic strength is used as a weight reflecting presynaptic selectivity.

## Methods

The code for simulations and figures is available at https://github.com/juliavg/decoding.

### Neuron model

#### Output neuron

The output neuron is modeled as a rate based neuron. Its firing rate *y* at time *t* is given by the equation:

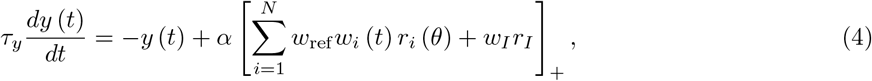

where *τ_y_* = 1 ms is the rate time constant, *α* = 0.1 nA^−1^ is the slope of the neuron’s transfer function, *w_ref_* = 16 nA is a reference weight for the *N* excitatory inputs, *w_i_* and *r_i_* are respectively the weight and firing rate of input neuron *i*, *w_I_* = –1700 pA and *r_I_* = 100 Hz are respectively the weight and firing rate of the inhibitory source. The inhibitory source could be understood as a single neuron firing at 100 Hz or multiple neurons firing at a lower rate and adding up to the same value. The sign []_+_ indicates a rectification that sets all negative values to 0.

#### Input neurons

The input neurons are modeled as rate based neurons with von Mises tuning curves, which is a “bump” like curve with circular boundary conditions. The firing rate of input neuron *i* at time *t* is given by:

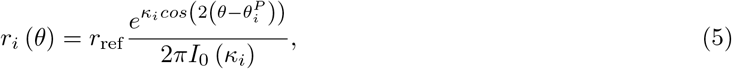

where *r_ref_* = 125 Hz is a reference firing rate, *θ* is the stimulus being shown to neuron *i* at time *t*, *κ_i_* is a parameter defining the width of the tuning curve from neuron *i*, 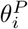 is the preferred orientation of neuron *i* and *I*_0_ is the modified Bessel function of order 0.

### Plasticity models

#### Variance (Figures 1–3)

The weights from the excitatory inputs are plastic, and evolve according to:

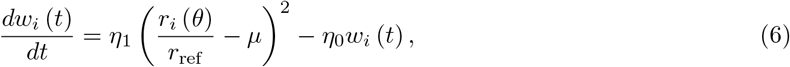

where *η*_1_ = 0.1 and *η*_0_ = 0.03 are learning rates and 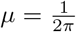 is a constant representing the mean activity of the presynaptic neurons.

#### Covariance (Figure S2)

For the simulations with the covariance rule, the excitatory weights evolve according to:

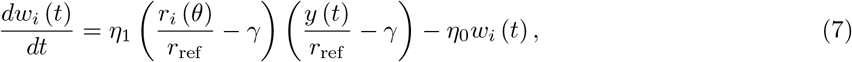

where *γ* = 0.24 is a constant and the remaining parameters are the same as with the rule based on presynaptic variance.

### Plasticity simulation (Figures 1 - 3)

One output neuron receives input from *N* = 50 neurons. Each input neuron *i* has a tuning curve defined by its width *κ_i_*, which is drawn randomly and independently for each neuron from a uniform distribution]0, 1], and by its preferred orientation 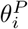, which is drawn randomly and independently for each neuron from a uniform distribution 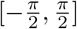.

The firing rate of the input neurons are initially set to *r_i_* = 20 Hz for a warm up period of 200 s, after which the stimulation protocol starts. During the stimulation protocol, a new orientation *θ* is chosen randomly from a uniform distribution 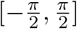 every *T* = 200 ms. The same orientation *θ* is shown to all input neurons and the whole stimulation protocol consists of 1 000 stimuli. We run 100 independent simulation runs, and shown in the figures is the pooled data from all of them.

### Orientation selectivity

Preferred orientation and orientation selectivity of neuron *i* are calculated from the circular mean of neuronal response *R_i_*:

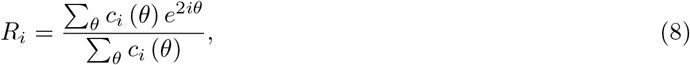

where *c_i_* (*θ*) is the calculated tuning curve of neuron *i*. We calculate *c_i_* (*θ*) as the mean response across stimuli using 20 bins on the interval 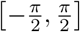 and using data from the last 500 stimuli in the simulation. Preferred orientation 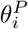 and the selectivity of neuron *i* are then calculated as the angle and the length of the resultant *R_i_*, respectively.

### Plasticity weights

The equilibrium weights for the plasticity rule based on presynaptic variance can be calculated by setting the left hand side of Equation 6 to zero:

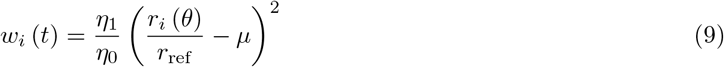

Assuming that the weights reach the steady state *W* within the interval *T* (how long each orientation is shown), we substitute *θ*(*t*) by the random variable Θ and take expected values:

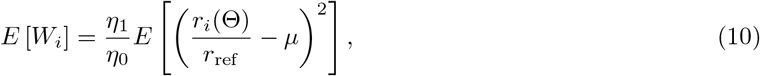

Finding 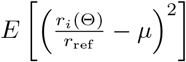, and substituting it in Equation 10 gives the equilibrium weight from presynaptic neuron *i*:

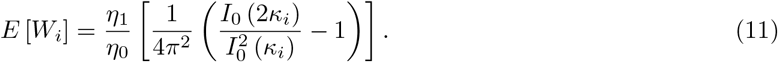

### Active input analysis (Figure 3)

For the active input analysis, we compare the mean activity of each input neuron during the presentation of a single stimulus to a threshold 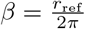 and consider it to be active (not active) when the activity is above (below) *β*.

### Maximum likelihood decoder

From Abbott and Dayan 2005, the estimated stimulus 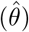 of a decoder based on maximum likelihood inference (ML decoder) can be determined by:

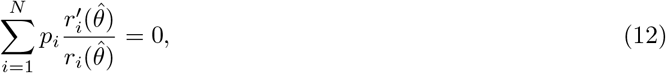

where *p_i_* is the firing rate of neuron *i*, *r_i_* (*θ*) is the tuning curve of neuron *i*, and the prime denotes derivative. Assuming the tuning curves of the simulation input neurons (Equation 5):

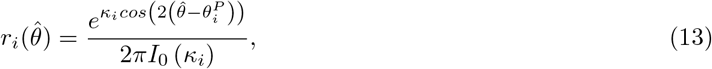

then the estimated stimulus 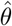 can be determined by:

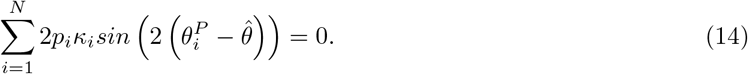

Solving for 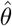 gives:

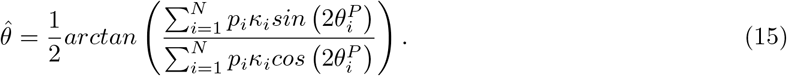

See also Keemink et al. 2018 for a similar derivation.

### Decoder simulation (Figure 4)

A decoder infers the stimulus *θ* shown to a population of 50 input neurons. The activity of input neuron *i* in response to stimulus *θ* (*p_i_*) is given as a sample randomly drawn from the Poisson distribution:

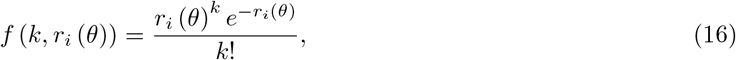

where *f*(*k, r_i_* (*θ*)) describes the probability of neuron *i* firing with rate *k* in response to stimulus *θ*, and *r_i_* (*θ*) is the tuning curve of neuron *i* (Equation 5).

The decoder estimates the shown stimulus according to:

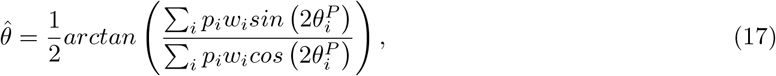

where *p_i_* is the firing rate of neuron *i* (obtained from sampling from Equation 16), *w_i_* are the weights used by the decoder and 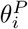 is the preferred orientation of neuron *i*. The tuning curves and parameters of input neurons are taken from the 100 independent plasticity simulations.

We compare the performance of 5 different decoders, which differ based on the weights *w_i_* they use: (i) *variance*: the weights used are obtained from the corresponding plasticity simulation; (ii) *ML*: *w_i_* = *κ_i_*; (iii) *uniform*: *w_i_* = 1; (iv) *shuffled*: the weights are the same as the *variance*, but shuffled; (v) *covariance*: the weights used are obtained from the plasticity simulations using the covariance plasticity rule.

For each independent plasticity simulation, we evaluate the decoder by showing 20 orientations equally spaced in the interval 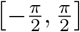. Each orientation is shown repeatedly for 100 trials, and we calculate the bias *best*(*θ*), variance 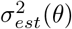 and error *e_est_*(*θ*) using:

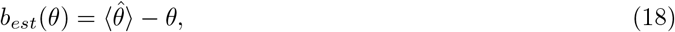

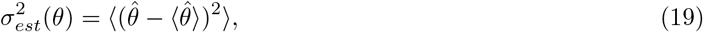

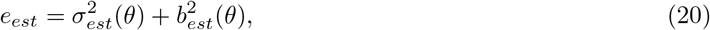

where 〈〉 indicates the average across trails. For each independent plasticity simulation, we average the bias *b_est_*(*θ*), variance 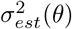 and error *e_est_*(*θ*) across stimuli *θ* to obtain a single value per simulation.

## Supplementary Material

**Supplementary Figure 1:**
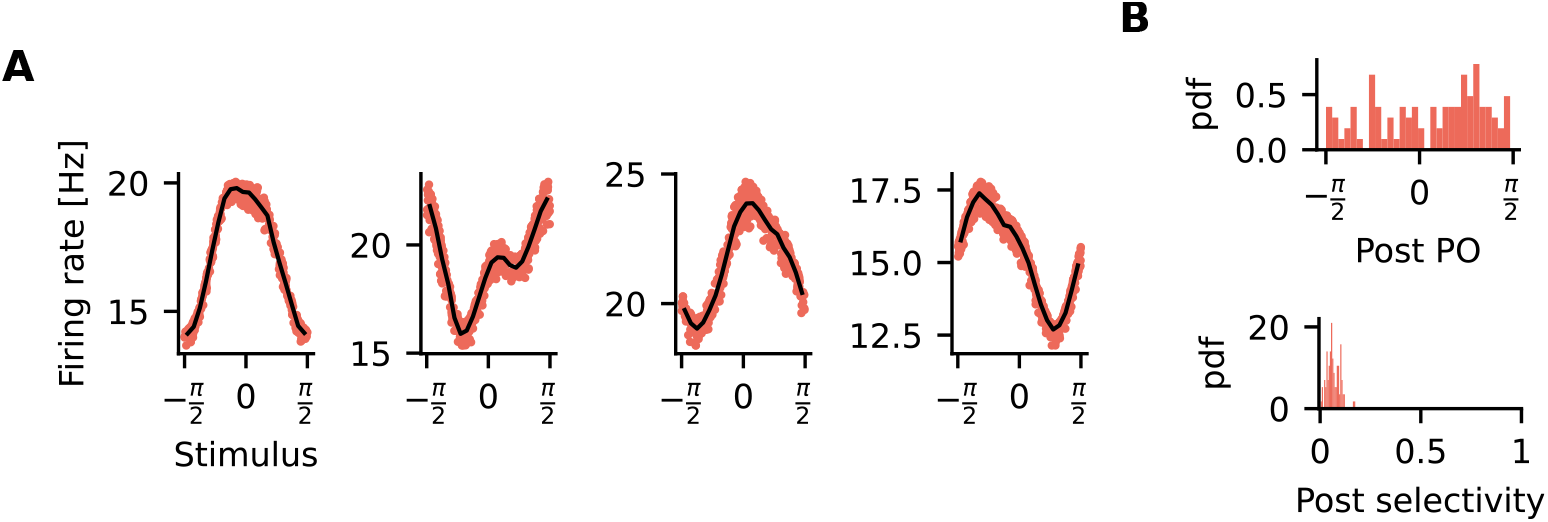
The effect of weaker inhibition on the selectivity of the postsynaptic neuron. The simulations performed for this figure are the same as those performed for Figure 2, except that the weight from the inhibiory source is 5 times weaker. (A) Examples of tuning curves for the postsynaptic neuron from 4 independent simulation runs. (B) Statistics of postsynaptic selectivity. Data pooled from 100 independent simulation runs. *Top*: Probability density function (pdf) of the postsynaptic PO. *Bottom*: Probability density function (pdf) of the postsynaptic selectivity.

**Supplementary Figure 2:**
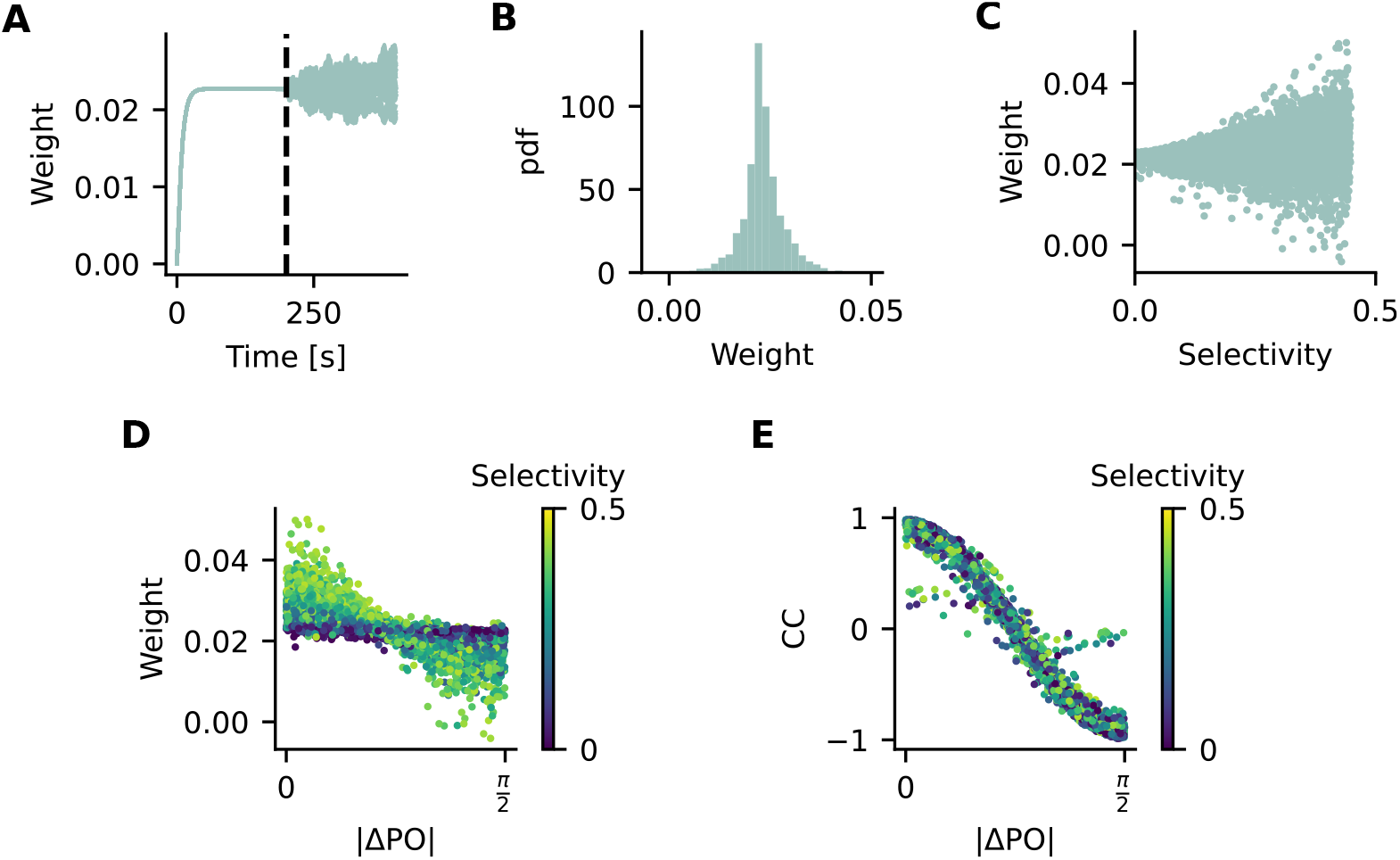
Relationship between synaptic weights and selectivity of presynaptic inputs in a simulation using the covariance plasticity rule. The simulations performed for this figure are the same as those performed for Figures 1–2, except that the plasticity rule used is the covariance rule (see Methods for details). (A) Synaptic weights as a function of time. Each line shows the synaptic weight of a single presynaptic neuron. The vertical dotted line indicates the moment when the stimulation protocol starts. (B) Distribution of synaptic weights at the end of the stimulation protocol. (C) Synaptic weights as a function of presynaptic selectivity. (D) Synaptic weights as a function of ΔPO. (E) Pearson’s correlation coefficient between pre- and postsynaptic neurons as a function of ΔPO. (D-E) Colors indicate presynaptic selectivity.

**Supplementary Figure 3:**
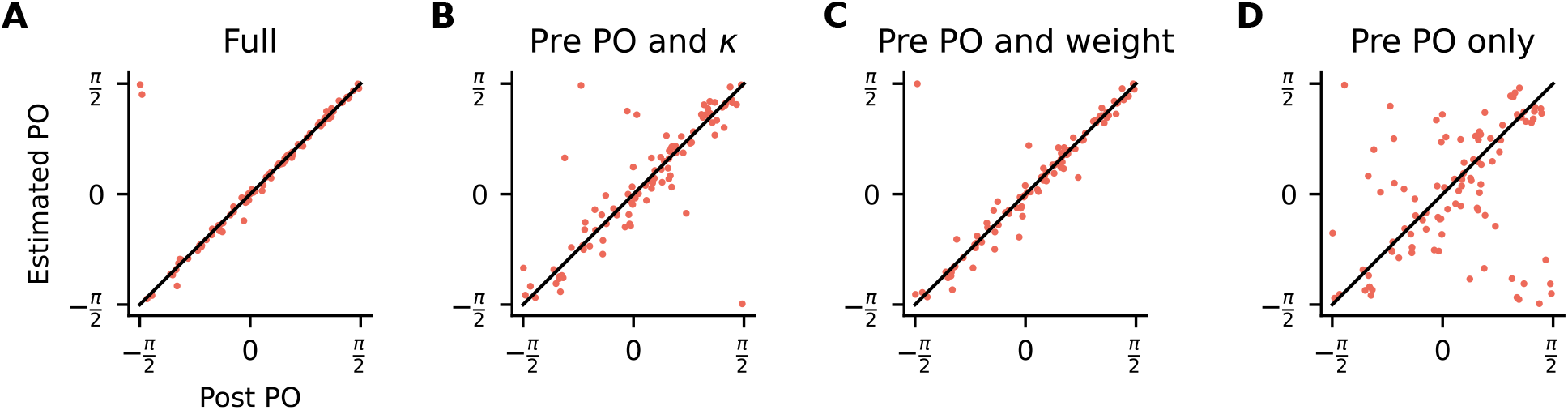
Postsynaptic PO is influenced by presynaptic PO, presynaptic selectivity and synaptic weights. (A) The postsynaptic tuning curve is estimated by adding the presynaptic tuning curves multiplied by the corresponding synaptic weights. The estimated postsynaptic PO is then extracted from the estimated postsynaptic tuning curve. (B) Same as in (A) but all weights are considered to be equal *w_i_* = 1. (C) Same as in (A) but all presynaptic tuning curves are considered to have the same width *κ_i_* = 1. (D) Same as in (A) but with *w_i_* = 1 and *κ_i_* = 1.

